# Prior Knowledge Biases the Visual Memory of Body Postures

**DOI:** 10.1101/2022.12.01.518647

**Authors:** Qiu Han, Marco Gandolfo, Marius V. Peelen

## Abstract

Body postures provide information about others’ actions, intentions, and emotional states. Little is known about how postures are represented in the brain’s visual system. Considering our extensive visual and motor experience with body postures, we hypothesized that priors derived from this experience may systematically bias visual body posture representations. We examined two priors: gravity and biomechanical constraints. Gravity pushes lifted body parts downwards, while biomechanical constraints limit the range of possible postures (e.g., an arm raised far behind the head cannot go down further). Across three experiments (N=246) we probed participants’ visual memory of briefly presented postures using change discrimination and adjustment tasks. Results showed that lifted arms were misremembered as lower and as more similar to the nearest biomechanically plausible postures. Inverting the body stimuli eliminated both biases, ruling out visual confounds. These findings show that visual memory representations of body postures are modulated by a combination of category-general and category-specific priors.

## Introduction

Body posture is an important social cue that provides information about others’ emotions, intentions, and mental states. The pressure to quickly and accurately recognize bodies and their movements has resulted in humans’ typically excellent performance in detecting and discriminating body posture and body motion ^1–4^, a skill that is supported by dedicated brain regions in visual cortex including the extrastriate body area ^5^, fusiform body area ^6^, and superior temporal sulcus ^7^. When bodies are presented inverted, which is inconsistent with our daily experience, the ability to detect and discriminate postures is impaired ^2,3,8–10^. This inversion effect is more pronounced for faces and bodies than for other objects, indicating more configural processing for these visually highly familiar stimuli ^2,9^.

Owning a body ourselves, we also have extensive motor, tactile, and proprioceptive experience of a body and its dynamics ^11^. Neuropsychological evidence suggests that we have an internal model of the physical relationships between body parts that helps us execute our own actions and understand those of others ^12^. Together with our extensive visual experience, these sensory modalities provide us with additional knowledge of hierarchical limb structure, the possible range of movements of joints, and the effort required for executing specific body actions. Here, we asked whether our experience observing and executing a biased range of body postures modulates the perceptual representation of these postures.

Previous research has shown that perception is influenced by knowledge and expectations ^13,14^. Specifically, Bayesian accounts of perception propose that priors are integrated with sensory input, weighted by their uncertainty to support perceptual inference ^15–17^. An example of this integration is the hollow-face illusion: a mask viewed from the concave side still gives a vivid impression of a convex face, due to the strong prior of faces being convex. Priors are shaped by environmental statistics, including the distribution of visual properties like orientation ^18,19^, basic physical principles of motion ^20,21^, gravity ^22,23^, and physical state ^24^.

Effects of prior knowledge on perception have also been observed for the perception of body movements. For example, observers tend to perceive or imagine a biomechanically plausible movement compared to an awkward one ^25^. Furthermore, when observing apparent body movements, the perceived movement tends to follow a biomechanically plausible path, even if that path is longer ^26^. Other examples include the finding that the extrapolation of biomechanically plausible movements is larger than implausible ones ^27^, and that unstable postures leaning backward are judged to be more likely to fall than postures leaning forward ^28^. These findings indicate that the perceptual interpretation of real or apparent body movements is influenced by knowledge of biomechanical constraints. However, body movements involve sequences of postures unfolding over time, requiring the viewer to predict and construct the upcoming posture. A single static posture may not automatically evoke such predictive processes. It is therefore unclear whether perceptual representations of static postures are influenced by priors in the way that body movements are.

To address this question, we considered two priors that are relevant for static body postures. The first is the general prior of gravity: Gravity is an omnipresent force that pushes everything down, resulting in a strong prior for perception and action ^29^. Previous studies have found that the position of an unsupported object will be remembered as lower, in line with the influence of a gravity prior ^30,31^. Accordingly, because arms will fall if not supported by muscles, we hypothesized that a lifted arm will be remembered as slightly lower than its actual position.

The second prior follows from biomechanical constraints. Because of the biomechanical structure of the body, particularly the range of motion of joints, postures are confined to a limited range. For example, the shoulder joints can flex from the resting position to the front by 180 degrees, but to the back, they can only extend to around 60 degrees ^32,33^. If prior knowledge of these constraints informs perception, the representation of a nearly impossible posture may be biased towards the nearest possible posture. Crucially, biomechanical constraints can counteract the effect of gravity: an arm raised in front of the head will fall but an arm raised behind the head can hardly fall lower (Figure 1).

**Figure 1.**
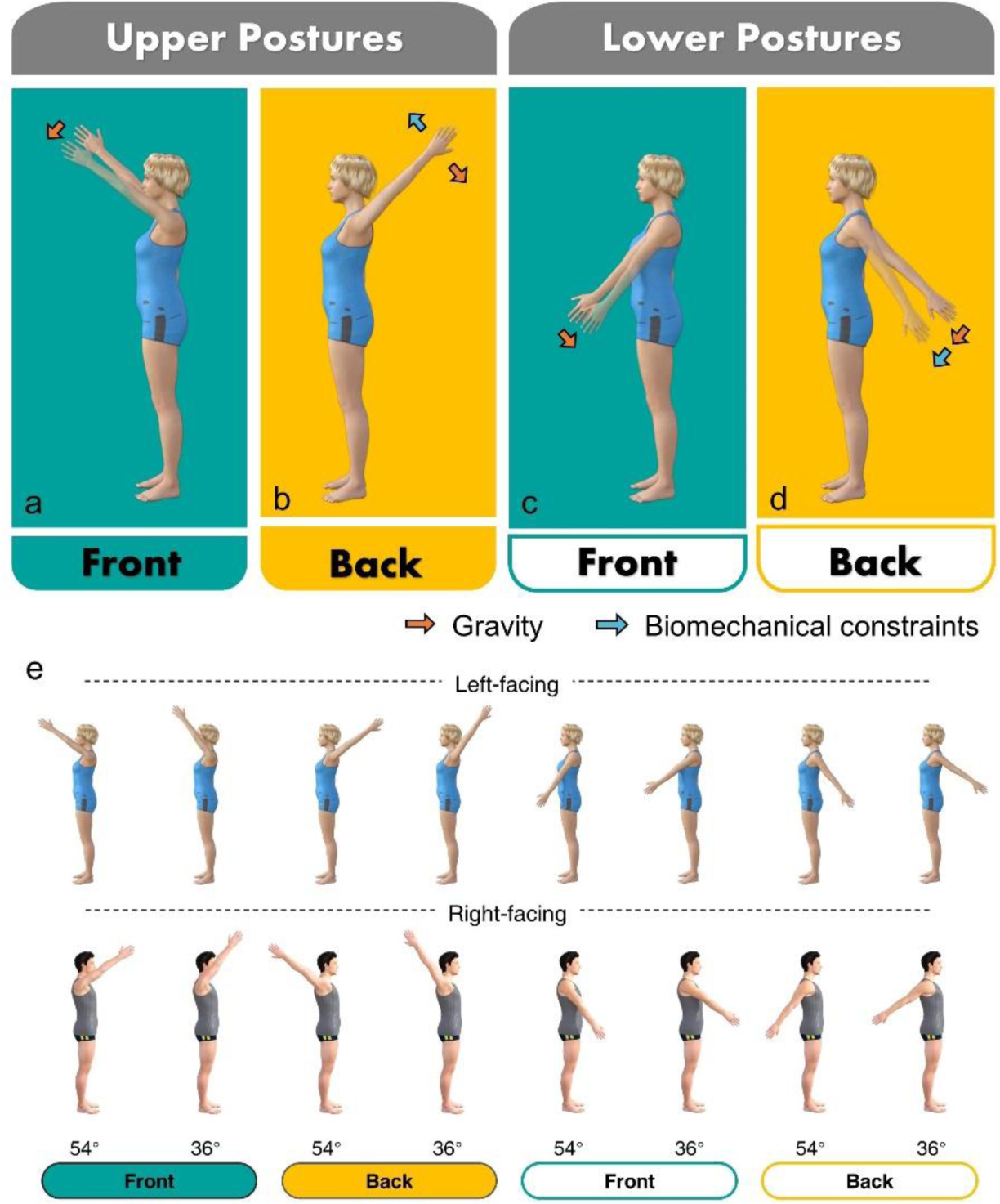
a)∼d), Illustration of the hypothesis. The orange and blue arrows indicate the direction of the gravity and the biomechanical constraints, respectively; transparent arms indicate the predicted perceived arm positions according the two hypotheses. Here the arrows and the predicted arms only suggest the direction but not the extent of the effects. For the Upper-back posture (b), gravity and the biomechanical constraints point in opposite direction, potentially eliminating each other’s influence. For the lower-back posture (d), gravity and the biomechanical constraints go in the same direction, their effects potentially adding up. e), Stimulus examples. For each quadrant, we show the lowest posture (54°) and the highest posture (36°) used in the experiments. In the experiment, both figures had left-facing and right-facing versions, though only one of them is shown here.

To test these hypotheses, we used four arm postures subject to one or both of the two biases (Figure 1). We predicted that lifted arms will generally be remembered as lower, towards the ground, reflecting a gravity-related bias. Furthermore, we predicted that lifted arms will be biased towards biomechanically possible postures. Specifically, biomechanical constraints limit further movement of the arm when the arm is raised behind the head, counteracting the gravity bias (Figure 1b), while adding to the gravity bias when the arm is behind the hip (Figure 1d).

We designed two tasks to probe the existence of biases in body posture representation due to gravity and biomechanical constraints. In the change discrimination task (Experiment 1, Figure 2a), participants compared two sequential postures whose arm positions slightly differed, with the second arm posture being slightly higher or lower. We first tested upper postures (Fig. 1a & 1b) in Experiment 1a, then followed with lower postures (Fig. 1c & 1d) in Experiment 1b to generalize the findings to visually different postures. We then replicated the results using a within-subject design with an adjustment task (Experiment 2, Figure 2b) where the participants needed to reproduce the remembered posture by adjusting the arm of a figure. The error of their adjustment reflects memory biases. Finally, we replicated these results again in Experiment 3, and used inverted body postures as control stimuli to test whether the effects rely on configural body processing.

**Figure 2.**
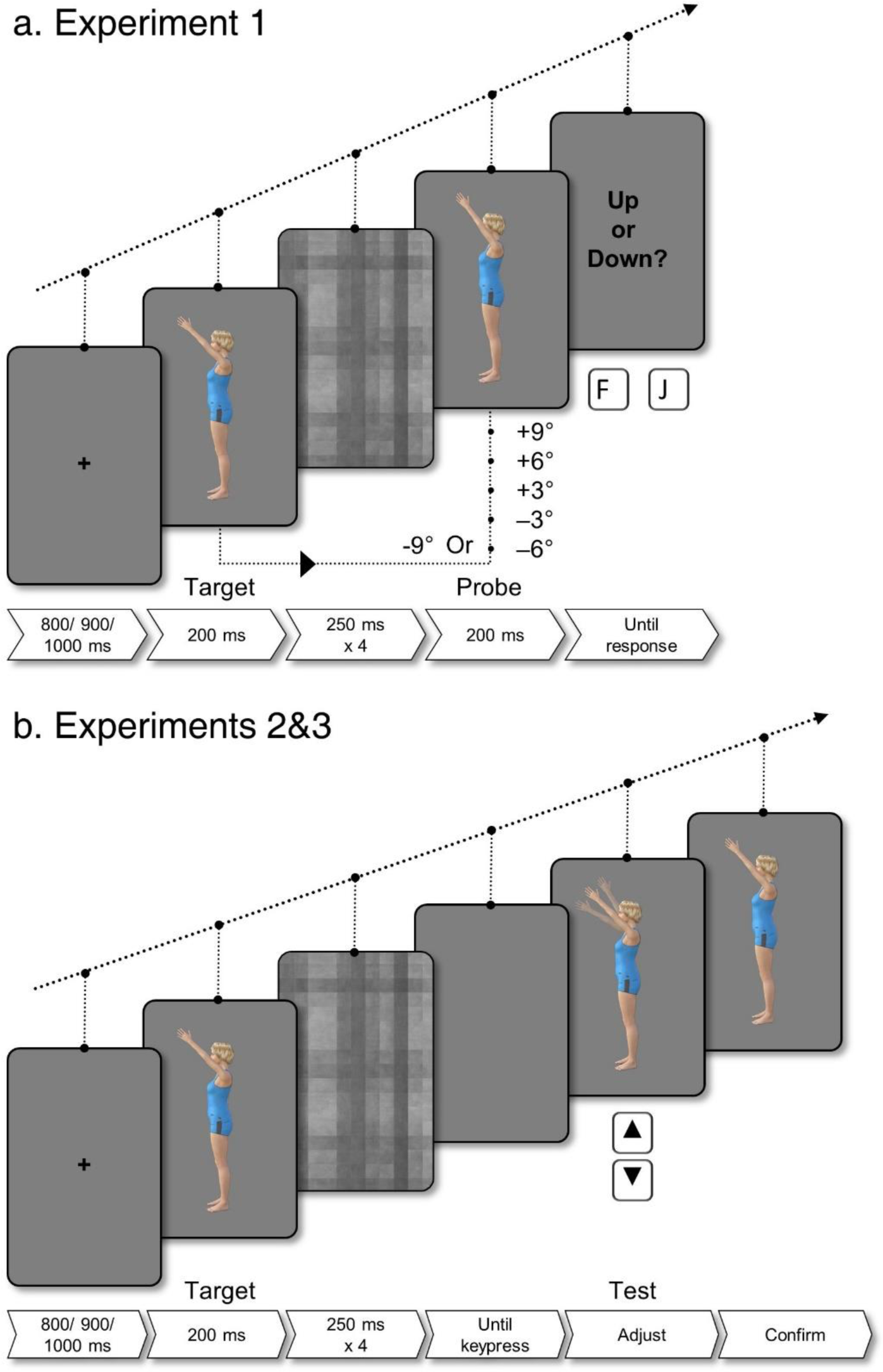
Trial Procedures. a) Change discrimination task used in Experiments 1a and 1b. In the trial shown here, the target is 45 degrees in the Upper-front, and the probe moves -9 degrees (i.e., upwards). Participants indicated whether the arm had moved up or down. The up-down text screen is shown for illustration purposes. b) Adjustment task used in Experiments 2 and 3. The target posture was either 36, 39, 42, 48, 51, or 54 degrees within each quadrant. The starting posture in the test image was chosen randomly from 30 to 60 from the same quadrant as the target. Participants adjusted the arm, indicated by transparent arms, using the up-arrow and down-arrow keys to match the target angle.

## Results

### Experiment 1a & 1b

In this change discrimination task, participants needed to decide whether the arm in the second posture was higher or lower than in the first one (Figure 2a). In the absence of biases, participants should detect upward and downward changes equally well. Instead, if priors bias the representation of the first posture during the brief interval, we may observe that detecting a change in one direction is easier than a change in the other. The perceptual biases of interest were thus quantified by the criterion (c) from signal detection theory. Taking upward movement as the signal, a negative criterion means that participants responded more up than down. Data of Experiment 1a (upper postures, N = 60) and 1b (lower postures, N = 60) were pooled in the analysis.

Participants’ responses were in line with a gravity bias (Figure 3): The arm in the target posture was remembered as lower than its actual position, as indexed by a criterion significantly below zero for all postures (Upper-front: M = -0.25, 95% CI = [-0.32, -0.18], *t*(59) = -7.08, *p* < .001, *d* = -0.91, BF_10_ = 5.75E6; Upper-back: M = -0.144, 95% CI = [-0.21, -0.07], *t*(59) = -4.38, *p* < .001, *d* = -0.57, BF_10_= 413; Lower-back: M = -0.11, 95% CI = [-0.19, -0.04], *t*(59) = -2.96, *p* = .004, *d* = -0.38, BF_10_ = 7.14) but not the Lower-front: M = -0.06, 95% CI = [-0.14, 0.02], *t*(59) = -1.60, *p* = .114, *d* = -0.21, BF_10_ = 0.47.

**Figure 3.**
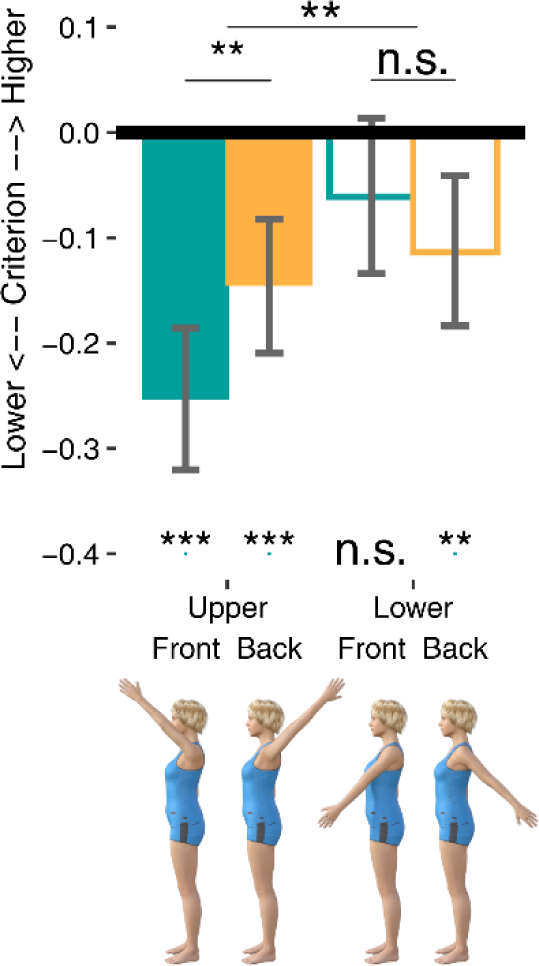
Criterion results for the four conditions in Experiment 1. A negative criterion reflects a bias to respond “up”, indicating that the first posture was remembered as lower than the second posture. We interpret this overall bias as reflecting knowledge of gravity. The difference between Front and Back indicates that the criterion was influenced by whether the arm is at an extreme posture. Results showed an interaction between Front/Back and Upper/Lower, in line with biomechanical constraints (see Figure 1). ***: p < .001, **: p < .01, *: p < .05, n.s.: not significant. Error bars denote 95% CI.

Next, we combined Experiments 1a and 1b using a mixed ANOVA with arm height (upper, Experiment 1a; versus lower, Experiment 1b) as a between-subject factor and arm direction as a within-subject factor to test the presence of a biomechanical bias. As illustrated in Figure 1, compared to the front, the downward bias in the back should be diminished by biomechanical constraints when in the upper quadrant (Figure 1a vs. 1b), but strengthened when in the lower quadrant (Figure 1c vs. 1d). We thus predicted an interaction between arm height and arm direction. We indeed found this interaction (Figure 3): *F*(1, 118) = 8.09, *p* = .005, η^2^_p_ = 0.064, BF_10_ = 7.63. Specifically, for the upper postures, the gravity bias was stronger in the front than in the back, M = -0.108, 95% CI = [-0.03, -0.19], *t*(59) = -2.74, *p* = .008, *d* = -0.35, BF_10_ = 4.17, indicating that the upward biomechanical constraint counteracted the gravity bias. The downward bias for the Lower-back was numerically stronger than that for the Lower-front, in line with an additive effect of biomechanical constraint bias and gravity bias, but this difference did not reach significance: M = 0.05, 95% CI = [-0.03, 0.13], *t*(59) = 1.3, *p* = .197, *d* = 0.17, BF_10_ = 0.32. All statistics are provided in Table S1, S2, and S3.

### Experiment 2

The change discrimination task provided evidence for both gravity and biomechanical biases. We wondered whether these results would be specific to the change detection task, in which the two consecutive body postures may be perceived as part of an action. If so, the results could reflect biases in human action perception rather than biases in the static representation of the target posture. To address this, in Experiment 2 (N=60) we tested whether the identified biases replicate in an adjustment task (Figure 2b), in which participants were asked to remember a target posture and then to reproduce this target posture by adjusting the arm of a human figure on the screen.

In the adjustment task, the direction and magnitude of biases are directly reflected in the direction and magnitude of the adjustment error. A negative error indicates that the target was remembered as lower than its actual position, reflecting a gravity bias. This was the case for all of the postures tested (Figure 4a, Upper-front: M = -2.54, 95% CI = [-2.97, -2.11], *t*(59) = -11.8, *p* < .001, *d* = -1.52, BF_10_ = 1.68E14; Upper-back: M = -1.89, 95% CI = [-2.34, -1.44], *t*(59) = -8.46, *p* < .001, *d* = -1.09, BF_10_ = 9.86E8; Lower-back: M = -1.07, 95% CI = [-1.48, -0.66], *t*(59) = -5.22, *p* < .001, *d* = -0.67, BF_10_ = 6.52E3) except for the lower-front (M = 0.09, 95% CI = [-0.35, 0.52], *t*(59) = -0.39, *p* = .694, *d* = 0.05, BF_10_ = 0.15).

**Figure 4.**
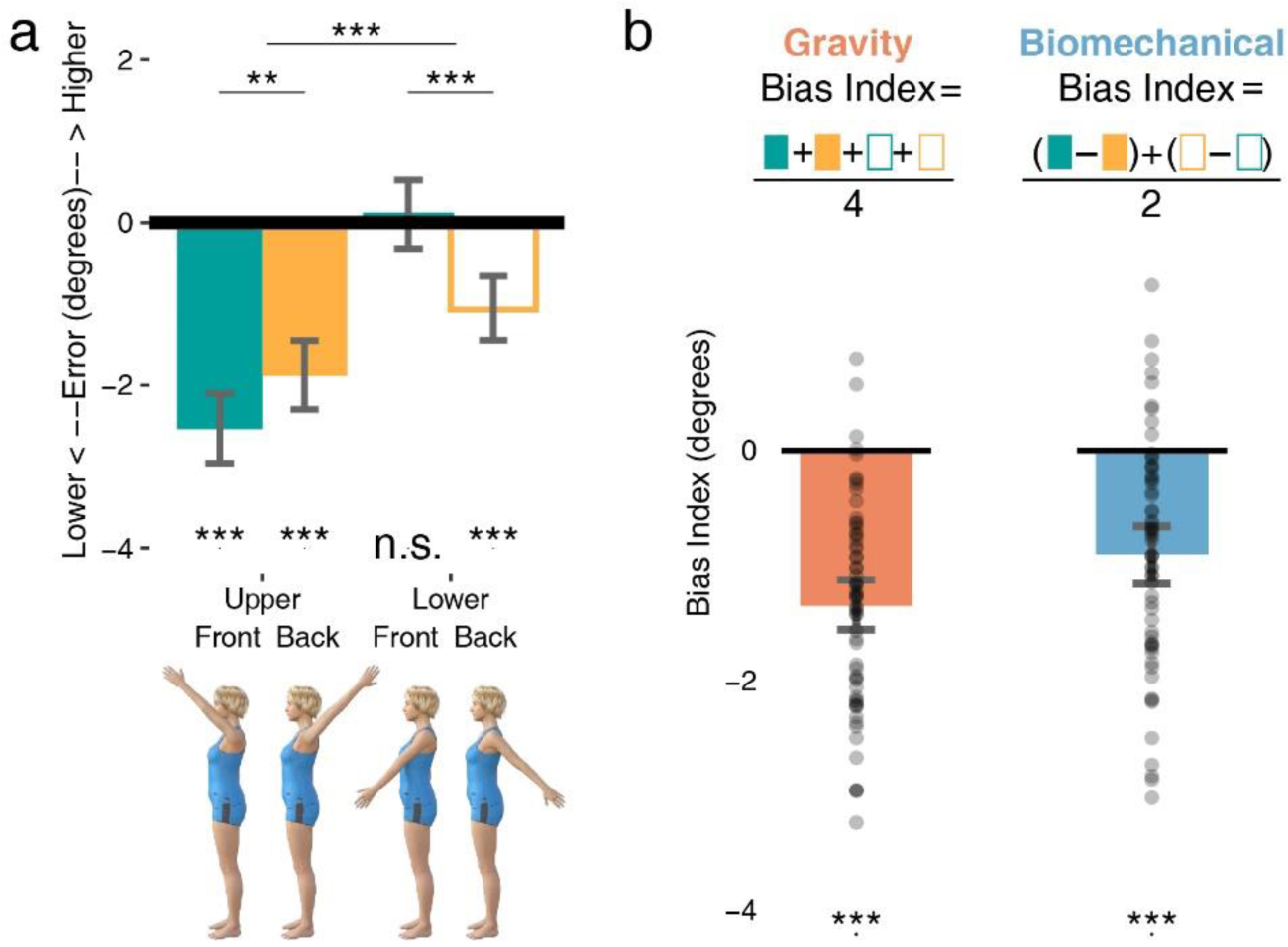
Results of Experiment 2. a) Mean error of the four conditions. b) Bias Indexes for individual participants. On the top are the calculation methods for the two indexes. ***: p < .001, **: p < .01, *: p < .05, n.s.: not significant. Error bars denote 95% CI.

Also consistent with Experiment 1, a two-way repeated-measures ANOVA showed an interaction between arm height and arm direction, revealing the effect of biomechanical constraints, *F*(1, 59) = 48.1, *p* < .001, η^2^_p_ = 0.45, BF_10_ = 2.02E8. As in Experiment 1, for the upper postures, the gravity bias was stronger in the front than the back: M = -0.65, 95% CI = [-0.25, -1.05], *t*(59) = -3.28, *p* = .002, *d* = -0.42, BF_10_ = 16.5. By contrast, as predicted, for the lower postures, the gravity bias was stronger in the back than the front: M = -1.15, 95% CI = [-0.79, -1.52], *t*(59) = -6.32, *p* < .001, *d* = -0.82, BF_10_ = 3.44E5, showing that biomechanical constraints also influence visual memory of lower arm postures.

For visualization purposes, we computed two indexes that reflect the two hypothesized effects. The gravity bias was given by the overall adjustment error, averaged across the four conditions (Upper-front, Upper-Back, Lower-front, Lower-back). The biomechanical constraint bias was indexed by the difference between postures with vs without biomechanical constraint, averaged across upper and lower postures (the mean of (Upper-front – Upper-back) and (Lower-back - Lower-front); Figure 4b). Figure 4b shows the bias indexes for individual participants. Taking the four postures together, the error caused by gravity was significantly different from zero, M = -1.35, 95% CI = [-1.58, -1.12], *t*(59) = -11.61, *p* < .001, *d* = -1.50, BF_10_ = 8.50E13. The overall biomechanical constraint (M = -0.90, 95% CI = [-1.16, -0.64], *t*(59) = -6.94, *p* < .001, *d* = -0.90, BF_10_ = 3.37E6, also reflected in the interaction in the ANOVA) was also highly consistent across individuals. These results confirm and extend the results of Experiment 1 using a different task, generalizing the effects to a scenario where no action or motion is implied.

### Experiment 3

Experiment 3 aimed to test whether the effects were caused by local visual features, including the overlap between the arm and the head, the curvature of the arm, and so on. A prominent feature of body perception is its susceptibility to inversion. Inversion has been shown to disrupt body and face perception more than other objects, which is believed to reflect the disrupted configural processing of bodies and faces ^2,3,8,10,34^. Inverted bodies thus serve as an ideal control for typical upright bodies since they are identical in terms of local features but are processed less as integral postures. In Experiment 3, we tested whether the effects of gravity and biomechanical constraints were contingent on the configural processing instead of local features using the inversion effect (N = 66).

We found a significant three-way interaction among arm direction, arm height and body orientation, *F*(1,65) = 7.15, *p* = .009, η^2^_p_ = .10, BF_10_ = 10.3, indicating that body orientation modulated the interaction between arm height and arm direction. Inspecting upright and inverted conditions separately (Figure 5), the interaction of arm height and arm direction was significant for the upright body, replicating results from Experiment 2, *F*(1,65) = 24.1, *p* < .001, η^2^_p_ = .27, BF_10_ = 8.50E4. In contrast, the inverted body did not show the interaction between arm height and arm direction, *F*(1,65) = 1.91, *p* = .171, η^2^_p_ = .029, BF_10_ = 0.55. A main effect of body orientation (*F*(1,65) = 46.9, *p* < .001, η^2^_p_ = .42, BF_10_ = 2.92E6) showed that inversion diminished the overall negative adjustment error, indicating a reduced gravity bias (Figure 5).

**Figure 5.**
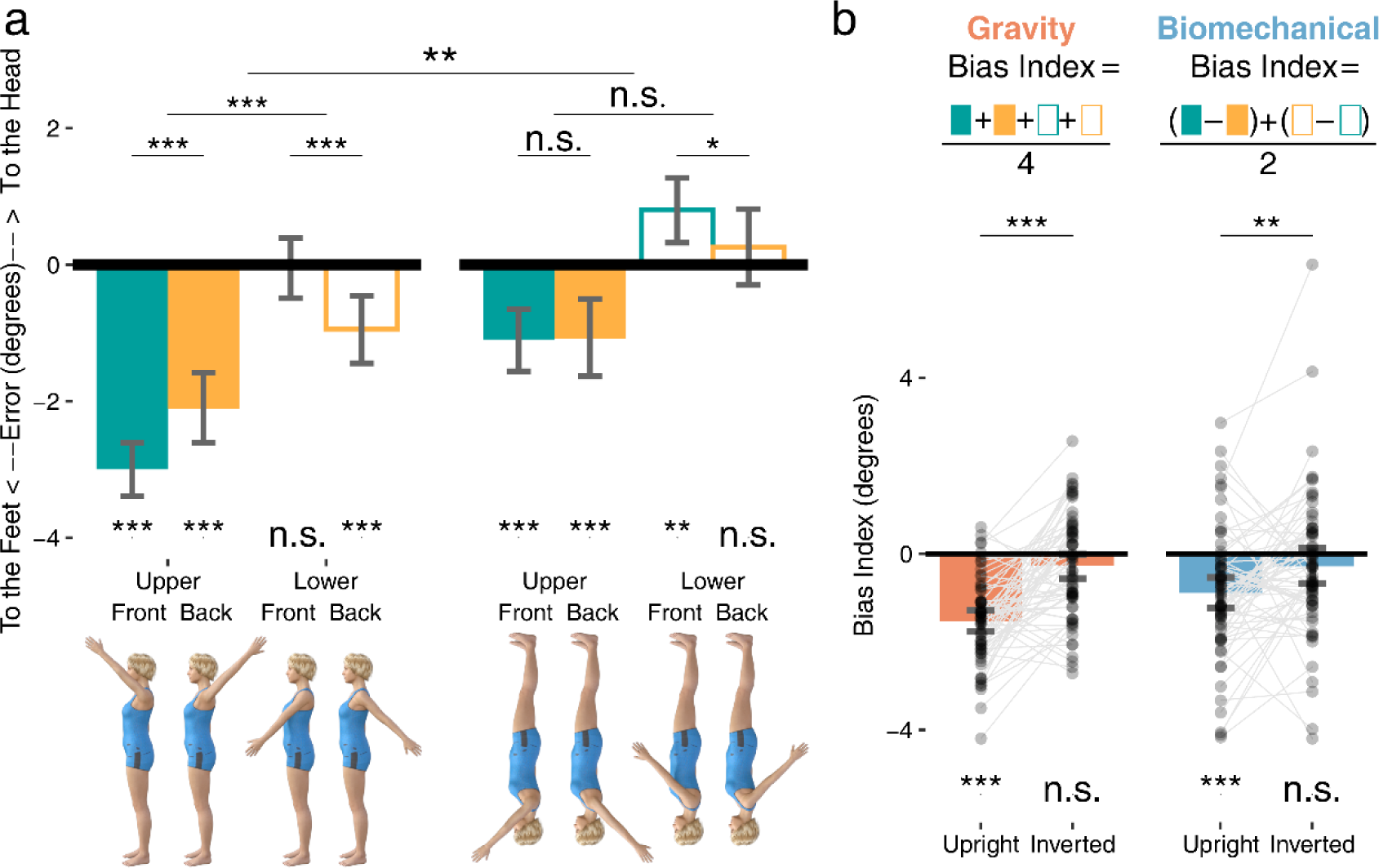
Results of Experiment 3. a) Mean error of the four conditions. Left: upright, Right: inverted. b) Bias Indexes for individual participants. On top are the calculation methods for the two indexes. ***: p < .001, **: p < .01, *: p < .05, n.s.: not significant. Error bars denote 95% CI.

As in Experiment 2, bias indexes were also calculated for visualization purposes and for a more direct description of the inversion effect. As shown in Figure 5b, inversion significantly reduced both gravity bias: M = 1.25, 95% CI = [0.88, 1.61], *t*(65) = 2.67, *p* < .001, *d* = 0.84, BF_10_ = 3.52E6 and biomechanical constraints: M = 0.61, 95% CI = [0.15, 1.06], *t*(65) = 2.67, *p* = .009, *d* = -0.33, BF_10_ = 3.54. This experiment excluded the possibility that these biases emerge from part-based processing of bodies or by the stimuli’s low-level visual features.

## Discussion

The current results demonstrate that priors resulting from gravity and biomechanical constraints jointly shape visual memory representations of human body postures. In three experiments, these effects were replicated both by directly repeating the same task and by using a different task. Importantly, the biases were absent when bodies were inverted, ruling out low-level visual confounds and indicating that the biases emerge from whole-body representations. Together, the two tasks we used excluded potential confounds of motion perception, response biases and local visual feature processing, demonstrating a top-down influence on static body representations.

Our results are well explained by Bayesian theories in which perception is the result of an interplay between sensory input and priors ^15,35^. Priors are shaped by environmental statistics ^18^ and serve to achieve optimal inference ^17^. In the case of body postures, multiple regularities jointly shape the prior distribution of postures, including biomechanical constraints that confine postures to a certain range and gravity that pulls limbs downward. When input is more ambiguous, priors will have a stronger influence, such that we tend to perceive what is most likely according to our prior. In our task, a body posture had to be maintained in visual memory for a brief interval. The fidelity of the sensory information will be reduced during this interval, making the posture representation susceptible to the influence of priors. Following this account, the biases should become larger when the uncertainty about the stimuli is increased, which can be tested in future studies, for example by degrading the stimulus or by increasing the memory interval.

Besides gravity and biomechanical constraints, the prior distribution of postures is also influenced by other factors. For example, common postures and movements like standing, walking, and using tools will lead to a relatively high probability of arms being in the lower front position. This might also explain the current finding that the lower front arm postures were judged and adjusted more accurately than the other postures (see Figure 3, 4a, 5a). Analyzing posture probabilities in large video databases and measuring biases at a higher resolution of the posture space would help establishing the link between posture probability and the biases observed here.

An alternative interpretation of the current findings is that they reflect visuomotor simulation, where observed postures are simulated in the viewer’s motor system as a mechanism to understand postures ^36^. This interpretation has been used to account for the finding of smaller representational momentum for biomechanically awkward arm movements ^37^. However, although the motor system has been shown to be activated during action observation ^38,39^, motor simulation may not be essential for understanding actions or representing body postures, as individuals born without upper limbs exhibit similar performance in action observation, action prediction, and mental imagery of postures ^40,41^. Knowing whether the effects observed here are also present in individuals born without limbs will give us more insight into the contributions of visual and motor experience. Furthermore, if motor experience plays a role in the visual memory of body postures, we might also expect an influence of commonly self-experienced postures ^42,43^. Accordingly, factors that influence an individual’s motor experience, including age ^44^, obesity ^45^, pain ^46,47^, and other clinical conditions, could potentially influence the biases reported here.

The current findings also raise new questions about the neural representation of body postures. Neuroimaging research on body perception has provided evidence for multiple cortical areas that are specifically engaged in body perception, including the extrastriate body area (EBA, Downing et al., 2001) and the fusiform body area (FBA, Peelen & Downing, 2005; Schwarzlose et al., 2005). Compared to the more extensively studied fusiform face area (FFA, Kanwisher et al., 1997), little is known about the representational structure of these areas ^51^. It has been shown that the FFA represents faces in a face space centered around the average face, with distances from the center representing the deviation from the mean face ^52,53^. Based on the current results that knowledge of body structure informs posture perception, the body-selective areas may store an internal model of the body, including its constraints. Given the many combinations of body part postures, it would be advantageous for neurons to be tuned primarily to biomechanically possible postures. Indeed, previous work has shown that body representations in the EBA more strongly represent postures in commonly experienced visual field locations ^54^. Our results suggest that the representational space of body postures in body-selective regions might be biased, reflecting perceived rather than physical distances between postures.

In sum, we show that body posture representation is biased towards the ground and towards biomechanically plausible postures, indicating an influence of both general knowledge of the world and specific knowledge of the body. These findings may reflect the influence of an internal model of the body based on environmental statistics. By employing such an encoding scheme, the visual system can efficiently predict upcoming postures, a critical component for humans’ ability to read others’ actions, intentions, and social interactions ^55^.

### Limitations of the study

This study revealed that the visual memory of a lifted-arm posture is modulated by knowledge about the world and the biomechanics of the body. A limitation of our study is that we only tested arm postures; future studies need to generalize our findings to other postures, for example leg postures. Another limitation is that our study could not address the origin of the prior. Future research is needed to determine whether visual experience, motor experience, or both contribute to the biases found here.

## STAR★Methods

### RESOURCE AVAILABILITY

#### Lead contact

Further information and requests for resources and reagents should be directed to and will be fulfilled by the lead contact, Qiu Han.

#### Materials availability

All the stimuli generated for this study are publicly available at https://osf.io/qmtkw/.

#### Data and code availability

All the codes, and raw data related to the experiments reported here are publicly available at https://osf.io/qmtkw/.

Additional information required to reanalyze the data reported in this paper is available from the lead contact upon request.

### EXPERIMENTAL MODEL AND STUDY PARTICIPANTDETAILS

All the studies were conducted on online platforms. Participants of Experiment 1 were recruited using the SONA system in return for course credits. Participants of Experiments 2 and 3 were recruited through Prolific in return for monetary reward. We required the participants to be above 18 years old with normal or corrected-to-normal vision. Digital informed consent was obtained from all participants. The procedures were approved by the University’s Ethical committee (Ethics no.: ECSW-2022-079).

The desired sample size was set to 60 for all experiments before testing. This was determined by power analysis using Jamovi suggesting that a sample size of 52 was needed to detect a minimum effect size of *d* = 0.4 with 80% power, as suggested recently as a first estimate for a reproducible effect size ^56^. We rounded this recommendation up to 60, resulting in 80% power to detect a minimum effect size of *d* = 0.368. Recruitment stopped when the sample size reached 60 after the exclusion of low-quality data (see METHOD DETAILS). For Experiment 1a, we recruited 75 participants. Of these, one participant did not finish the task and 14 were excluded. For Experiment 1b, we recruited 67 participants. Of these, two did not finish the task and five were excluded. We thus acquired an effective sample size of 60 for both Experiment 1a (49 females, 11 males; age: M = 20.1, range = [18, 36]) and Experiment 1b (48 females, 12 males, 2 other; age: M = 20.6, range = [18, 47]).

Experiment 2 adopted a within-subject design. The sample size was kept consistent with Experiment 1. 60 subjects (30 females, 30 males; age: M = 33.42, range = [21, 45]) were recruited and no one was excluded.

In Experiment 3, 66 participants (21 females, 45 males; age: M = 30.04, range = [18, 45]) were recruited. First, we recruited the intended sample size of 60, however, the number of participants starting with the upright vs inverted condition was not yet balanced when the sample size reached 60, therefore six additional participants were recruited. Data from only the first 60 participants yielded highly similar results.

## METHOD DETAILS

### Experiment 1

**Stimuli**: Body images were generated by rendering digital human models in DAZ studio 4.15 (Daz Productions, Inc). A female character and a male character were used. The characters were standing in profile, one arm lifted, the other leaning naturally on the hip. The lifted arm positions were categorized into four quadrants (Figure 1): two directions (front and back) x two arm heights (upper and lower). In each quadrant, we designated the upper bound of that quadrant as the zero point, with larger angles meaning that the arm is lower, i.e., closer to the feet. The arm could be presented at an angle of 36, 39, 42, 45, 48, 51, or 54 degrees in each quadrant. Both left-facing and right-facing figures were generated, so that an arm in front of the body was equally often presented in the left and right visual field to avoid possible confounds of visual field differences between the conditions (Figure 1e). The lifted arm was always on the viewer’s side (right arm lifted when facing right, left arm lifted when facing left) to avoid the arm being occluded by other body parts.

Mask images were grey-scale checkerboard images. Body images were 300 pixels wide, 480 pixels high. Size in degree depended on the online participant’s screen resolution and the eye-to-screen distance. Mask images were 350 pixels wide, 525 pixels high. Masks were presented slightly larger than the body to achieve a better masking effect.

### Procedures

Experimental procedures were programmed with jsPsych library ^57^ and the psychophysics plugin ^58^. Experiments 1a and 1b tested the front and the back arm directions for upper postures (Experiment 1a) and lower postures (Experiment 1b). Data were aggregated for analysis. Experiment 1a also included a machine condition which was not relevant to the purpose of the current study (see Figure S2).

In each trial, a fixation cross was first presented at the center for 800, 900, or 1000 ms, then a target body posture of either 36, 39, 42, 45, 48, 51, or 54 degrees in either quadrant was shown for 200 ms (Figure 2a). Participants were instructed to remember the posture of the target and hold it in memory. Immediately after the target, a 1000-ms dynamic mask consisting of four consecutive checkerboard images (250 ms each) was shown to minimize aftereffects and/or apparent motion of the arm, after which the probe image appeared for 200 ms. Compared to the target, the arm in the probe would move upwards or downwards by an angle of 3, 6, or 9 degrees (equiprobable). The task was to judge whether the arm had moved up or down relative to the target. Participants indicated their choice by pressing F or J on the keyboard. The key-response mapping was counterbalanced across participants. Participants were asked to respond as accurately and as quickly as possible. The trial ended upon response or 4000 ms after the probe had disappeared.

Each combination of arm direction and angle difference included 24 trials, resulting in 288 trials in total, separated into four blocks. Angle difference, arm direction, and figure gender were completely interleaved while facing direction was kept identical within blocks to avoid extra effort for switching viewpoint between trials. The two left-facing and two right-facing blocks were in ABBA order, with about half of the subjects starting with a left-facing block and the others starting with a right-facing block. A practice session of 12 trials was delivered before the formal experiment. Feedback on the accuracy and mean response time across conditions were shown to the participant at the end of each block.

### Experiment 2

Experiment 2 included the same four conditions as in Experiment 1. In each quadrant, six target angles (36, 39, 42, 48, 51, and 54) were used. Consistent with Experiment 1, a fixation and then the target posture was shown, followed by the mask. After the mask disappeared, the subjects were instructed to press one of the left-arrow or right-arrow keys to show the test image for adjustment. The initial posture of the test was randomized between 30 and 60 degrees, but always in the same quadrant as the target. Participants then pressed up and down arrow keys to manipulate the arm of the test image to move upwards or downwards. After adjusting the arm to the remembered target position, participants pressed space to confirm their answer. If no response was made, the trial ended after 10 s. If the test image was not initiated within 3 s after the mask, the trial skipped and participants were warned to start the adjustment more quickly in the following trials.

All the other factors, were kept consistent with Experiment 1. Facing direction was blocked, and other factors were interleaved. Each angle in each quadrant was presented eight times, resulting in 48 trials for each quadrant, 192 trials in total. The trials were divided into four blocks, with the order manipulated as in Experiment 1. Two mini practice blocks were completed before the start of the experiment. Feedback on average absolute error was given at the end of each block.

### Experiment 3

The task was identical to Experiment 2 except that both upright and inverted conditions were tested. The inverted body images were generated by vertically flipping the upright images. Half of the participants started with the upright condition and half with the inverted condition. Both upright and inverted conditions contained a left-facing block and a right-facing block, with the order randomized across participants but kept the same for upright and inverted conditions. Within each block, trials of different combinations of arm direction, arm height, within-quadrant angle, and figure identities were interleaved. Because of the inclusion of the inverted condition, the trial number for each angle was halved compared to Experiment 2. In total, Upper-front, Upper-back, Lower-front, and Lower-back all included 24 trials for the upright and 24 for the inverted condition.

### QUANTIFICATION AND STATISTICAL ANALYSIS

#### Experiment 1

Responses with RTs < 250 ms (relative to probe onset) were excluded (0.10% of the total number of trials across Exp 1a and 1b), as these most likely reflect anticipatory responses. For each participant, data quality was inspected by plotting the percentage of up responses for each angle difference level in each quadrant (Figure S1a). Participants following task instructions should show an increase in the percentage of up responses as the angle difference decreases. That is, the more obvious that the arm moves up, the more likely people choose up, yielding a sigmoid curve. Some participants exhibited a flat or reversed curve, suggesting that they misunderstood the task or pressed randomly. These participants were detected using a slope index (Figure S1b) of the difference between the mean up response percentage of the two most obvious moving-up levels (-9 and -6) and the mean of the two most obvious moving-down levels (9 and 6). Participants with a slope index below 0.2 in either the front or the back condition were excluded, resulting in 14 exclusions in Experiment 1a and 5 exclusions in Experiment 1b.

We calculated individual criterion in each condition from the hit rate (up response percentage when the arm actually moved up) and false alarm rate (up response percentage when the arm actually moved down) using the Psycho package ^59^ in R ^60^:

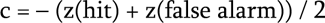

Statistics were done using bruceR package in R and JASP (JASP Team, 2023). Both frequentist results and Bayesian factor (BF) were provided. By convention, a BF (the ratio of the probability of acquiring the data given one model against another) > 3 indicates some evidence supporting the first model ^61^. For BF ANOVA, each effect was tested by comparing models that contain that effect against their equivalent models stripped of the effect. Results of d prime were also analyzed and are presented in Figure S1. For simplicity, statistics of effects of interest are reported in the Results section and all the other statistics are provided in Table S1, S2, and S3.

#### Experiment 2

Trials in which participants pressed space before adjusting the test image and trials in which the test image was not initiated within 3 s after the mask were discarded. Trials with an absolute error larger than 15 degrees were also discarded. Altogether, this led to the rejection of 5.41% of the total number of trials. No participants were excluded.

The error of the response from the target was used as the index of biases, calculated as the target angle minus the response angle. For example, a target of 48 degrees adjusted as 52 degrees would yield an error of -4 degrees. Negative values indicate that the posture was adjusted to be lower than it was actually shown. Errors of all trials in one quadrant were averaged to get the mean error for each quadrant of each participant. Group mean error data were tested in the same way as the criterion in Experiment 1, except that a repeated-measures ANOVA was used instead of a mixed ANOVA.

#### Experiment 3

The coding of angles for the inverted condition followed a body-centered reference frame rather than a spatial reference frame. For example, negative errors for an upright body indicate that the arm was adjusted as closer to the feet and thus closer to the lower part of the screen. Similarly, negative errors for an inverted body indicate that the arm was adjusted as closer to the feet; however, because of the inversion and the reference to the body, this is now closer to the upper part of the screen.

Data cleaning procedures were identical to Experiment 2, with an exclusion rate of 6.90% of the total number of trials. Adjustment errors for each quadrant were averaged for upright and inverted conditions separately. A three-way repeated-measures ANOVA with arm height, arm direction, and body orientation was conducted. Two bias indexes for upright and inverted conditions were calculated and compared with two-tailed t-tests to test whether inversion diminished the biases.

## Acknowledgments

We thank Paul E. Downing and Eelke Spaak for feedback on earlier versions of the manuscript and the Peelen Lab members for the helpful suggestions on experimental design during lab meetings.

This project has received funding from the China Scholarship Council (CSC), European Union’s Horizon 2020 research and innovation programme under the Marie Skłodowska-Curie (grant agreement No. 101033489), and European Research Council (ERC) under the European Union’s Horizon 2020 research and innovation programme (grant agreement No 725970).

## Author Contributions

Qiu Han: Conceptualization, Methodology, Software, Investigation, Formal Analysis, Validation, Visualization, Writing – Original Draft, Writing – Review & Editing, Funding Acquisition

Marco Gandolfo: Conceptualization, Methodology, Software, Validation, Writing – Review & Editing, Funding Acquisition

Marius Peelen: Conceptualization, Methodology, Validation, Writing – Review & Editing, Funding Acquisition

