## Supplementary Figures and Tables for "Prior Knowledge Biases the Visual Memory of Body Postures"

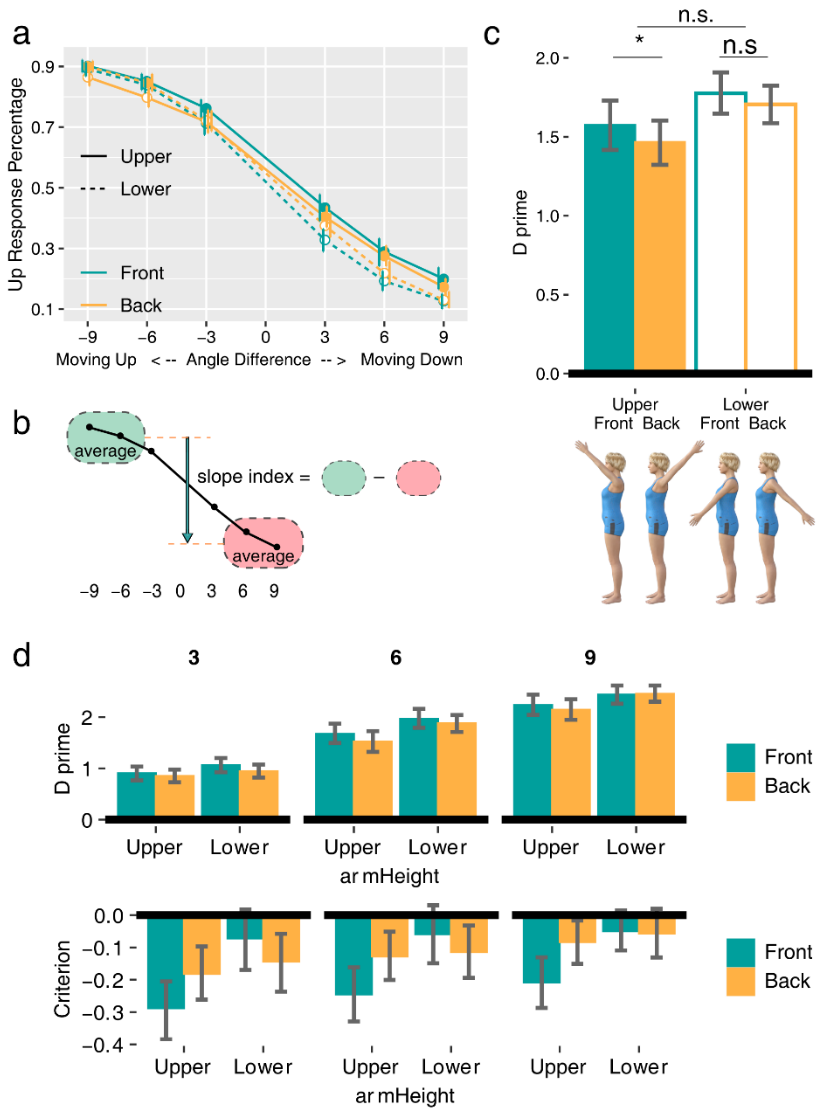


**Figure S1. Additional Results of Experiment 1**, related to Figure 3. a: Results of up response percentage against angle difference for four postures. b: illustration of slope index calculation used in Experiment 1 for data exclusion (see STAR method). The black solid curve represents an up response percentage pattern of an individual participant. A flat curve corresponds to a lack of discrimination between up and down trials and a small slope index. This slope index was thus used to exclude participants who did not perform the task according to the instructions. c. Results of d prime (sensitivity index). Sensitivity was generally higher for lower than upper postures: $F(1, 118) = 5.45$, $p = .021$, $\eta^2_p = 0.04$, and higher for front than back: $F(1, 118) = 6.52$, $p = .012$, $\eta^2_p = 0.05$. These effects might be related to familiarity with different postures in daily experience: arms in the lower positions and front of the body are more frequently seen and performed. The interaction between arm height and arm direction, however, was absent for d prime: $F(1, 118) = 5.45$, $p = .58$, $\eta^2_p = 0.003$. d. Results of d prime and criterion broken down by angle differences (3, 6, 9 degrees). For criterion, the interaction between arm height and arm direction is visible across all angle differences.

*: $p < .05$, n.s.: not significant. Error bars denote 95% CI.

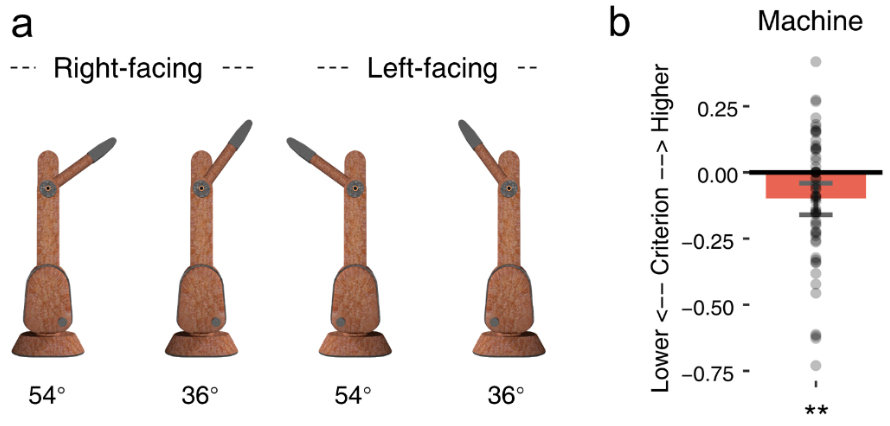


**Figure S2. Stimuli and Results of the Machine Condition**, related to Figure 3. a. the machine stimuli. A machine with a mechanical arm was created using SketchUp (Trimble Inc.). The height of the main body and of the joint linking the mechanical arm and the main body were kept consistent with the human body. We did not distinguish the front and back sides of the machine, thus the machine had only one upper "posture" instead of both Upper-front and Upper-back postures as for the body. Images were mirrored to keep the arm in both left and right view fields. Consistent with the body, 36, 39, 42, 45, 48, 51, and 54 degrees were used. Here only the highest and lowest positions are shown.

b. Results of the machine condition. This experiment used the same 75 participants as Experiment 1a. Based on a slope index (Figure S1c) calculated on the Up response percentage of the machine, sixteen participants were excluded, resulting in a sample size of 59 (49 females, 10 males; age: M = 20.5, range = [18, 36]). Half of the participants started with the machine blocks (144 trials) and the other half with the body blocks. The change discrimination task was identical to Experiment 1. We found a bias towards the lower direction for the machine: M = -0.10, 95% CI = [-0.16, -0.04], $t(58) = -3.16$, $p = .003$, $d = -0.41$, BF10 = 11.8. This result aligns with previous studies reporting a gravity bias for unsupported objects [S1, S2]. However, it is also possible that participants interpreted the machine as a human body, similar to a cartoon character. This single experiment thus does not allow for general conclusions about a possible gravity bias for inanimate objects.

**: $p < .01$. Error bar denotes 95% CI.

**Table S1. Statistics of Experiment 1**, related to Figure 3.

**a.** ANOVA (Mixed Design), output from bruceR package in R

Descriptives:

| "armHeight" | "direction" | Mean | S.D. | n |
|---|---|---|---|---|
| Upper | Front | -0.252 | (0.276) | 60 |
| Upper | Back | -0.144 | (0.255) | 60 |
| Lower | Front | -0.061 | (0.294) | 60 |
| Lower | Back | -0.114 | (0.298) | 60 |

Total sample size: *N* = 120

ANOVA Table:
Dependent variable(s): c
Between-subjects factor(s): armHeight
Within-subjects factor(s): direction
Covariate(s): –

| | MS | MSE | df1 | df2 | F | p | | η²p [90% CI of η²p] | η²G |
|---|---|---|---|---|---|---|---|---|---|
| armHeight | 0.734 | 0.110 | 1.000 | 118.000 | 6.661 | .011 | * | .053 [.007, .132] | .038 |
| direction | 0.046 | 0.048 | 1.000 | 118.000 | 0.951 | .331 | | .008 [.000, .055] | .002 |
| armHeight * direction | 0.388 | 0.048 | 1.000 | 118.000 | 8.094 | .005 | ** | .064 [.011, .147] | .020 |

**b.** Bayes Factor, output from JASP.

| Effects | P(incl) | P(excl) | P(incl\|data) | P(excl\|data) | BFincl |
|---|---|---|---|---|---|
| direction | 0.400 | 0.400 | 0.087 | 0.376 | 0.232 |
| armHeight | 0.400 | 0.400 | 0.367 | 0.096 | 3.827 |
| direction ✻ armHeight | 0.200 | 0.200 | 0.537 | 0.070 | 7.634 |

*Note.* Compares models that contain the effect to equivalent models stripped of the effect. Higher-order interactions are excluded. Analysis suggested by Sebastiaan Mathôt.

**Table S2. Statistics of Experiment 2**, related to Figure 4.

**a.** ANOVA (Within-Subjects Design), output from bruceR package in R

Descriptives:

| "direction" | "armHeight" | Mean | S.D. | n |
|---|---|---|---|---|
| Front | Upper | -2.540 | (1.666) | 60 |
| Front | Lower | 0.085 | (1.668) | 60 |
| Back | Upper | -1.890 | (1.731) | 60 |
| Back | Lower | -1.070 | (1.587) | 60 |

Total sample size: *N* = 60

ANOVA Table:
Dependent variable(s): Error
Between-subjects factor(s): –
Within-subjects factor(s): direction, armHeight
Covariate(s): –

| | MS | MSE | df1 | df2 | F | p | | $\eta^2 p$ [90% CI of $\eta^2 p$] | $\eta^2 G$ |
|---|---|---|---|---|---|---|---|---|---|
| direction | 3.830 | 1.161 | 1.000 | 59.000 | 3.299 | .074 | . | .053 [.000, .170] | .006 |
| armHeight | 178.009 | 5.632 | 1.000 | 59.000 | 31.608 | <.001 | *** | .349 [.192, .485] | .214 |
| direction * armHeight | 48.849 | 1.015 | 1.000 | 59.000 | 48.125 | <.001 | *** | .449 [.294, .571] | .070 |

**b.** Bayes Factor, output from JASP.

| Effects | P(incl) | P(excl) | P(incl\|data) | P(excl\|data) | BFincl |
|---|---|---|---|---|---|
| armHeight | 0.400 | 0.400 | 1.671×10-8 | 6.021×10-13 | 27758.736 |
| direction | 0.400 | 0.400 | 4.941×10-9 | 1.177×10-8 | 0.420 |
| armHeight ✻ direction | 0.200 | 0.200 | 1.000 | 4.941×10-9 | 2.024×10+8 |

*Note.* Compares models that contain the effect to equivalent models stripped of the effect. Higher-order interactions are excluded. Analysis suggested by Sebastiaan Mathôt.

**Table S3. Statistics of Experiment 3**, related to Figure 5.

**a.** ANOVA (Within-Subjects Design), output from bruceR package in R

Descriptives:

| "direction" | "armHeight" | "inversion" | Mean | S.D. | n |
|---|---|---|---|---|---|
| Front | Upper | Upright | -2.997 | (1.730) | 66 |
| Front | Upper | Inverted | -1.095 | (1.815) | 66 |
| Front | Lower | Upright | -0.058 | (1.911) | 66 |
| Front | Lower | Inverted | 0.807 | (1.989) | 66 |
| Back | Upper | Upright | -2.104 | (2.256) | 66 |
| Back | Upper | Inverted | -1.081 | (2.373) | 66 |
| Back | Lower | Upright | -0.942 | (2.116) | 66 |
| Back | Lower | Inverted | 0.256 | (2.378) | 66 |

Total sample size: *N* = 66

ANOVA Table:
Dependent variable(s): Error
Between-subjects factor(s): –
Within-subjects factor(s): direction, armHeight, inversion
Covariate(s): –

| | MS | MSE | df1 | df2 | F | p | | η²p [90% CI of η²p] | η²G |
|---|---|---|---|---|---|---|---|---|---|
| direction | 2.294 | 2.421 | 1.000 | 65.000 | 0.948 | .334 | | .014 [.000, .095] | .001 |
| armHeight | 444.613 | 11.885 | 1.000 | 65.000 | 37.409 | <.001 | *** | .365 [.216, .494] | .164 |
| inversion | 205.229 | 4.380 | 1.000 | 65.000 | 46.857 | <.001 | *** | .419 [.270, .540] | .083 |
| direction * armHeight | 45.182 | 3.201 | 1.000 | 65.000 | 14.113 | <.001 | *** | .178 [.059, .316] | .020 |
| direction * inversion | 2.459 | 1.595 | 1.000 | 65.000 | 1.541 | .219 | | .023 [.000, .114] | .001 |
| armHeight * inversion | 6.135 | 4.227 | 1.000 | 65.000 | 1.451 | .233 | | .022 [.000, .111] | .003 |
| direction * armHeight * inversion | 12.136 | 1.697 | 1.000 | 65.000 | 7.152 | .009 | ** | .099 [.014, .226] | .005 |

**b.** Bayes Factor, output from JASP.

| Effects | P(incl) | P(excl) | P(incl\|data) | P(excl\|data) | BFincl |
|---|---|---|---|---|---|
| inversion | 0.263 | 0.263 | 0.344 | 1.179×10-7 | 2.921×10+6 |
| dire | 0.263 | 0.263 | 0.006 | 0.036 | 0.166 |
| armHeight | 0.263 | 0.263 | 0.030 | 1.335×10-7 | 227487.219 |
| inversion ✻ dire | 0.263 | 0.263 | 0.122 | 0.450 | 0.271 |
| inversion ✻ armHeight | 0.263 | 0.263 | 0.180 | 0.428 | 0.421 |
| dire ✻ armHeight | 0.263 | 0.263 | 0.564 | 0.008 | 74.121 |
| inversion ✻ dire ✻ armHeight | 0.053 | 0.053 | 0.392 | 0.038 | 10.289 |

*Note.* Compares models that contain the effect to equivalent models stripped of the effect. Higher-order interactions are exclude d. Analysis suggested by Sebastiaan Mathôt.
